# Estimating the statistical performance of different approaches to meta-analysis of data from animal studies in identifying the impact of aspects of study design

**DOI:** 10.1101/256776

**Authors:** Qianying Wang, Jing Liao, Kaitlyn Hair, Alexandra Bannach-Brown, Zsanett Bahor, Gillian L Currie, Sarah K McCann, David W Howells, Emily S Sena, Malcolm R Macleod

## Abstract

**Background:** Meta-analysis is increasingly used to summarise the findings identified in systematic reviews of animal studies modelling human disease. Such reviews typically identify a large number of individually small studies, testing efficacy under a variety of conditions. This leads to substantial heterogeneity, and identifying potential sources of this heterogeneity is an important function of such analyses. However, the statistical performance of different approaches (normalised compared with standardised mean difference estimates of effect size; stratified meta-analysis compared with meta-regression) is not known.

**Methods:** Using data from 3116 experiments in focal cerebral ischaemia to construct a linear model predicting observed improvement in outcome contingent on 25 independent variables. We used stochastic simulation to attribute these variables to simulated studies according to their prevalence. To ascertain the ability to detect an effect of a given variable we introduced in addition this “variable of interest” of given prevalence and effect. To establish any impact of a latent variable on the apparent influence of the variable of interest we also introduced a “latent confounding variable” with given prevalence and effect, and allowed the prevalence of the variable of interest to be different in the presence and absence of the latent variable.

**Results:** Generally, the normalised mean difference (NMD) approach had higher statistical power than the standardised mean difference (SMD) approach. Even when the effect size and the number of studies contributing to the meta-analysis was small, there was good statistical power to detect the overall effect, with a low false positive rate. For detecting an effect of the variable of interest, stratified meta-analysis was associated with a substantial false positive rate with NMD estimates of effect size, while using an SMD estimate of effect size had very low statistical power. Univariate and multivariable meta-regression performed substantially better, with low false positive rate for both NMD and SMD approaches; power was higher for NMD than for SMD. The presence or absence of a latent confounding variables only introduced an apparent effect of the variable of interest when there was substantial asymmetry in the prevalence of the variable of interest in the presence or absence of the confounding variable.

**Conclusions:** In meta-analysis of data from animal studies, NMD estimates of effect size should be used in preference to SMD estimates, and meta-regression should, where possible, be chosen over stratified meta-analysis. The power to detect the influence of the variable of interest depends on the effect of the variable of interest and its prevalence, but unless effects are very large adequate power is only achieved once at least 100 experiments are included in the meta-analysis.

## Introduction

Meta-analysis is increasingly used to summarise information from animal studies which relate to a particular question relevant to human health. In this context a fundamental question is not just whether a given intervention improves outcome in animal models (they usually do); but also what the characteristics of that efficacy are, which might help guide the application of those findings to for instance clinical trial design. For instance, the design of the EuroHYP-1 trial (1), in particular the depth and duration of cooling, was based in part on findings of a systematic review of animal experiments (2). A large number of factors might influence the effects observed, for instance the species, age and sex of the experimental animal; the mode of induction and the duration of the cerebral ischaemia; the intervention, its dose and route of delivery; the means by which the outcome was measured; and risks of bias in the contributing experiments which might be addressed by randomisation, blinding etc. It is important to know how well the different approaches to meta-analysis perform in identifying the impact of these factors, both to inform the analytical choice and to form a judgement of how likely it is that an important modifying effect has been identified. Since the number of studies contributing to a meta-analysis is not an issue of investigator choice but of the amount of primary research available, such power considerations are important both to design (one might decide that if fewer than a certain number of studies were identified, meta-analysis would not be performed) and to interpretation (the absence of a significant finding in a large meta-analysis would be more compelling than if the meta-analysis included a small number of studies).

Such meta-analyses can use different approaches to representing the effect size and to accounting for heterogeneity (3). For example, a change in the volume of brain tissue damaged can either be expressed as a proportion of the damage seen in untreated controls (the “normalised mean difference” approach, NMD); or the difference between groups can be expressed in units relating to the observed variance, on the basis that the population variance should be constant across studies (the “standardised mean difference” approach, SMD). One concern is that, when group size is small, the sampled variance will depart more from the underlying population variance, introducing a measurement error which will reduce the precision of the estimate of effect size. Indeed, we have previously shown that, in seeking evidence for small study effects such as publication bias, the use of SMD estimates of effect size leads to funnel plot asymmetry even in the absence of publication bias (4).

The influence of study characteristics (independent variables) on the observed effect size can be determined by two methods. In the first, partitioning of heterogeneity, the sum of the heterogeneity observed within groups defined by certain characteristics is subtracted from the total heterogeneity, with the residual interpreted as the heterogeneity explained by the categorisation, tested against the Chisquared distribution with n-1 degrees of freedom, where n is the number of categories. A potential problem is that, due to sampling, some effect sizes may appear to be very precise, and therefore given substantial weight in the meta-analysis. Since the heterogeneity is the sum of the weighted squared deviations from the fixed effects estimate, any categorisation which places an influential (highly weighted) study closer to the fixed effect estimate in that category than it is to the global estimate will “explain’ some of the observed heterogeneity; and since the fixed effect estimate is most sensitive to the most influential studies, this is likely to occur. The second approach to investigating differences between groups of studies is to seek to fit the findings to a meta-regression model, testing whether any of the characteristics are associated with regression coefficients significantly different from zero.

These considerations are relevant to meta-analysis of both clinical and preclinical (animal) data, but we have been concerned that because individual animal experiments are generally smaller than clinical trial cohorts, the error in sampling of variance may weaken the SMD approach to calculating effect sizes and the use of partitioning of heterogeneity to explore differences between groups of studies.

We therefore undertook simulation studies, based on models derived from our large number of systematic reviews of the focal cerebral ischaemia literature, to better understand the weaknesses and any strengths of these approaches. Given that meta-regression can involve different approaches to the estimation of tau, we also compared these approaches.

## Methods

### Development of model for simulation

We identified 3116 experiments in focal cerebral ischaemia involving 48119 animals, identified in the context of systematic reviews and curated in the CAMARADES dataset. As well as the mean and variance for control and treatment groups the dataset included ten discrete, two continuous and thirteen binary variables. For discrete and binary variables the proportion of experiments in each category was calculated. The two continuous variables (time of treatment administration and time of assessment) did not show any linear relationship with outcome (Supplementary Table 1.1), so we divided these into quintiles, took the median value for each group and considered these as discrete variables. For time of outcome assessment, 10% of studies measured outcome at less than 24 hours and 29% of studies measured outcome at 24 hours, and so the first two quintiles were collapsed into one. The 25 variables, including their type, possible values and corresponding proportion are listed in Table 1.

**Table 1:** Variables offered to meta-analysis

| **Variables** | **Levels** | **Proportion** |
| --- | --- | --- |
| **Drug group** | Other | 18% |
|  | Stem Cells | 18% |
|  | Growth factor | 16% |
|  | Thrombolytic | 12% |
|  | Antidepressant | 6% |
|  | NOS Inhibitor | 6% |
|  | Anti-Inflammatory | 5% |
|  | PPAR-gamma agonist | 5% |
|  | Antioxidant | 9% |
|  | Vitamin | 4% |
| **Animal** | Rat | 80% |
|  | Mouse | 14% |
|  | Other | 6% |
| **Sex** | Male | 82% |
|  | Unknown | 9% |
|  | Female | 7% |
|  | Both | 3% |
| **Type of Ischaemia** | Temporary | 57% |
|  | Permanent | 29% |
|  | Other | 14% |
| **Anaesthetic** | Halothane | 39% |
|  | Isoflurane | 17% |
|  | Unknown | 11% |
|  | Chloral Hydrate | 9% |
|  | Ketamine | 9% |
|  | Other | 15% |
| **Method of Induction of Injury** | Intraluminal filament | 52% |
|  | Electrocoagulation | 9% |
|  | Microvascular clip | 8% |
|  | Donor clot | 6% |
|  | Autologous embolism | 6% |
|  | Photochemical thrombosis | 4% |
|  | Cauterisation | 4% |
|  | Other | 13% |
| **Method of Quantification of Injury** | TTC | 46% |
|  | H & E | 22% |
|  | Cresyl Violet | 6% |
|  | Unknown | 5% |
|  | MRI | 5% |
|  | Other | 15% |
| **Ventilation** | Unknown | 64% |
|  | Spontaneous | 22% |
|  | Other | 13% |
| **Route of Drug Delivery** | IVenous | 42% |
|  | IPeritoneal | 20% |
|  | SubCut | 11% |
|  | ICerebVentricular | 8% |
|  | Stereotactic | 8% |
|  | Other | 10% |
| **Comorbidity** | None | 91% |
|  | Other | 9% |
| **Cotreatment in both Groups** | TRUE | 8% |
|  | FALSE | 92% |
| **Cotreatment in Treatment Group Only** | TRUE | 4% |
|  | FALSE | 96% |
| **Peer Review Publication** | TRUE | 98% |
|  | FALSE | 2% |
| **Control of Temperature** | TRUE | 76% |
|  | FALSE | 24% |
| **Monitoring of Physiological Variables** | TRUE | 52% |
|  | FALSE | 48% |
| **Random Allocation to Group** | TRUE | 48% |
|  | FALSE | 52% |
| **Blinded Induction of Ischaemia** | TRUE | 24% |
|  | FALSE | 76% |
| **Blinded Assessment of Outcome** | TRUE | 47% |
|  | FALSE | 53% |
| **Anaesthetic without marked intrinsic neuroprotective activity** | TRUE | 73% |
|  | FALSE | 27% |
| **Use of Comorbid Animals** | TRUE | 11% |
|  | FALSE | 89% |
| **Sample Size Calculation** | TRUE | 2% |
|  | FALSE | 98% |
| **Compliance with Animal Welfare Regulations** | TRUE | 72% |
|  | FALSE | 28% |
| **Statement of Potential Conflicts of Interest** | TRUE | 12% |
|  | FALSE | 88% |
| **Time of Administration (hrs)** | -2 | 20% |
|  | 0.17 | 20% |
|  | 2 | 20% |
|  | 6 | 20% |
|  | 24 | 20% |
| **Time of Assessment (hrs)** | 24 | 40% |
|  | 72 | 20% |
|  | 168 | 20% |
|  | 672 | 20% |

For each of 23 nominal variables with *m* categories, we created *m*-1 dummy variables to represent these categories. These 23 categorical variables are represented in our analysis as 54 dummy variables. We built a weighted linear regression model with 54 dummy variables and two continuous variables where the effect size (response variable) and weight of each study were calculated using the normalised mean difference method (3). The resulting regression coefficients are shown in Supplementary Table 1.1.

### Simulation of studies to be used in meta-analysis

The purpose of the simulation was to observe the statistical power to detect the influence of a variable of interest; and whether the influence of a latent (unobserved) confounding variable might be misconstrued as representing an effect of the variable of interest. For instance, the ambient noise in an animal house might be an important determinant of outcome, but may be unreported. If randomisation were more commonly performed by investigators who had quieter animal houses, the effect of animal house noise might be misconstrued as an effect of randomisation.

There are 10 parameters for the simulation: (i) the number of studies simulated in the meta-analysis; (ii) the effect of the variable of interest; (iii) the prevalence of the variable of interest when the confounding variable is also present; (iv) the prevalence of the variable of interest when the confounding variable is absent; (v) the effect of the confounding variable; (vi) the prevalence of the confounding variable; (vii) the effect size in the absence of the variable of interest and the confounding variable; (viii) the pooled standard deviation (ix) the significance threshold (set to 0.05); and (x) the method used in estimating tau.

The choice of these variables was informed by our prior meta-analyses. For instance, in systematic reviews in focal cerebral ischaemia models the number of included experiments ranges from very low (5) to over 300 (2;6). In such reviews the overall effect size reported is around 25%; that is, a reversal of the disease phenotype to 75% of the response seen in untreated controls. Detecting an effect of less than 10% (i.e. 0.1 NMD) might not be practical. We set the pooled standard deviation to 20% of the modelled infarct volume for control animals (i.e. 0.2 NMD), based on the variance observed in the studies used to construct the model. The impact of aspects of study design such as randomisation or blinding or the choice of anaesthetic can also be expressed in this way, and we consider that, while many such factors will have some importance, those which shifted the estimate of efficacy by 10% (0.1 NMD, absolute, not relative) would be important to detect. Because of concerns about the number of subjects per variable (7) we did not perform meta-regression unless the number of studies (k) was 70 or higher, and we set our default value of k to 70.

We modelled experiments with characteristics determined by stochastic simulation using random number generation from the frequency of independent variables included in the regression model. The size of experimental cohorts was again determined by stochastic simulation based on the observed distribution of sample sizes, with a simplification that group sizes in the treatment and control groups be equal. For ease of modelling we assumed equal variance across all studies. We then used that regression equation to calculate a predicted “true” effect size for each simulated study. We then sampled individual animals from a control group population with mean infarct size of 100, and from a treatment group with a mean infarct size determined by the regression equation according to the modelled characteristics.

For each single simulated study we calculated the observed means and standard deviations for the control and treatment groups and calculated both NMD and SMD estimates of effect size and variance.

For the NMD estimates of effect size and, separately, for the SMD estimates, we then combined the k studies using a random effects meta-analysis to provide a global estimate of efficacy, calculated the z-value, and recorded whether this was significantly different from the null using the significance threshold determined above (Table 2). Next, we performed stratified meta-analysis by partitioning the heterogeneity due to the presence or absence of the variable of interest, and recorded whether the heterogeneity explained by the partitioning reached the threshold for significance, using the chi squared test, determined above. Next we performed univariate meta-regression where the response variable was the effect size and the covariate was the variable of interest, and recorded whether the coefficient of the variable of interest was significantly different from zero. Finally, we performed multivariable meta-regression where the response variable was the effect size, and the covariates included the previous described variables along with the variable of interest, but not of the unknown (latent) confounding variable; and recorded whether the coefficient of the variable of interest was significantly different from zero.

**Table 2:** Parameters provided in the simulation

| Parameter | Default value |
| --- | --- |
| Number of studies (k) | 100 |
| Effect of Variable of Interest (Vol) | 10 (NMD 0.1) |
| Prevalence of Vol within group with CF | 0.4 |
| Prevalence of Vol within group without CF | 0.2 |
| Effect of confounding variable (CF) | 10 (NMD 0.1) |
| Prevalence of confounding variable | 0.4 |
| Base effect size | 10 (NMD 0.1) |
| Pooled standard deviation | 20 (NMD 0.2) |
| Heterogeneity estimator | Restricted maximum likelihood |
| Significance level | 0.05 |

To generate stable estimates of power and false positive rates we ran these simulations 1000 times for each of NMD and SMD effect size measures, and repeated this while varying baseline assumptions described in Table 2. For each set of 1000 simulations, for each analysis, we calculated the number of simulations in which a statistically significant effect was observed. Where we had modelled an effect, this gave us an estimate of the statistical power to detect that effect. Where we modelled that there was no effect, this gave us an estimate of the false positive rate. If the test was performing as desired, we would expect this to be close to the threshold of significance determined in the modelling. The process is illustrated in a flow diagram in Supplementary Figure 1.1. The R code for these simulations, which uses the *metafor* package, is available at GitHub (https://github.com/qianyingw/power-simulation). We set the random seed generator in R from 1 to 1000 in each replication to allow others to repeat our work using the same parameters.

We ran the simulations under the Windows 8.1 x64 system, with core i705500U, CPU 2.40 GHz and 2.39 GHz, RAM 8.00 GB. Interestingly, there was no major improvement when running the simulations on a high performance Linux compute infrastructure consisting of 7000 Intel^®^ Xeon^®^ cores with up to 3 TB of memory available per compute node, perhaps because we did not optimise our code for a parallel computing environment.

## Results

The simulations took several days to run. As well as being dependent on the number of studies in each meta-analysis and the number of simulations run, computation time was influenced by the choice of tau estimation, being longer for the iterative methods (maximum likelihood, restricted maximum likelihood, Empirical Bayes) and shorter for the other methods (DerSimonian-Laird, Hedges, Hunter-Schmidt or Sidik-Jonkman).

**Table 3:**
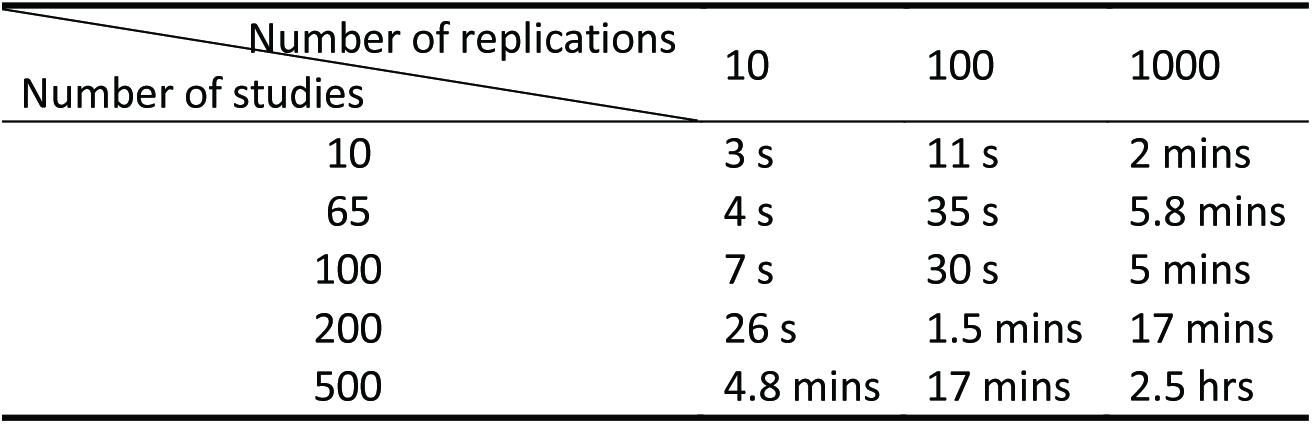
Performance of the simulation

For 1000 simulations of a meta-analysis involving 100 studies, we recorded the “true” (i.e. predicted) effect size in the absence of the variable of interest or the confounding variable and with the base effect size set to 0; this followed a normal distribution (Kolmogorov-Smirnov test, p=0.4526) with mean 0.0002 and standard deviation 2.383.

### Detecting overall effects and effects in subgroups

#### 1. Sensitivity to number of studies

We studied the power of the global meta-analysis to detect an effect size either when there was no effect present (blue in Fig 1a) or under the baseline assumptions of Table 2 (red in Fig 1a). The baseline true effect size is 16 (10 from the base assumption, 4 from the effect of the confounding variable (effect of 10 in 40% of studies), and 2.8 from the effect of the variable of interest (effect of 10 in 28% of studies). We varied the number of studies from 10 to 200. With no true effect present, the false positive rate remained at around 5%, as expected, for both NMD and SMD analyses. There was a difference between NMD and SMD when an effect was present, with 80% power to detect an effect seen with around 30 included studies for NMD, but requiring over 50 included studies for SMD.

In seeking to detect effects in subgroups, we first used partitioning of heterogeneity (Fig 1b). In combination with the use of an SMD estimate of effect size the false positive rate was constant at around 5%, and the power to detect the modelled difference in efficacy between subgroups increased as the number of studies increased. However, this power was only 31% with 100 studies in the meta-analysis and even with 200 studies was only around 49%. In contrast, the use of an NMD estimate of effect size was associated with a false positive rate which increased with the number of included studies: around 60% with 40 included studies and more than 80% with 200 included studies. The statistical power was only around 10% higher than the false positive rate across the range of study sizes studied.

**Figure 1:**
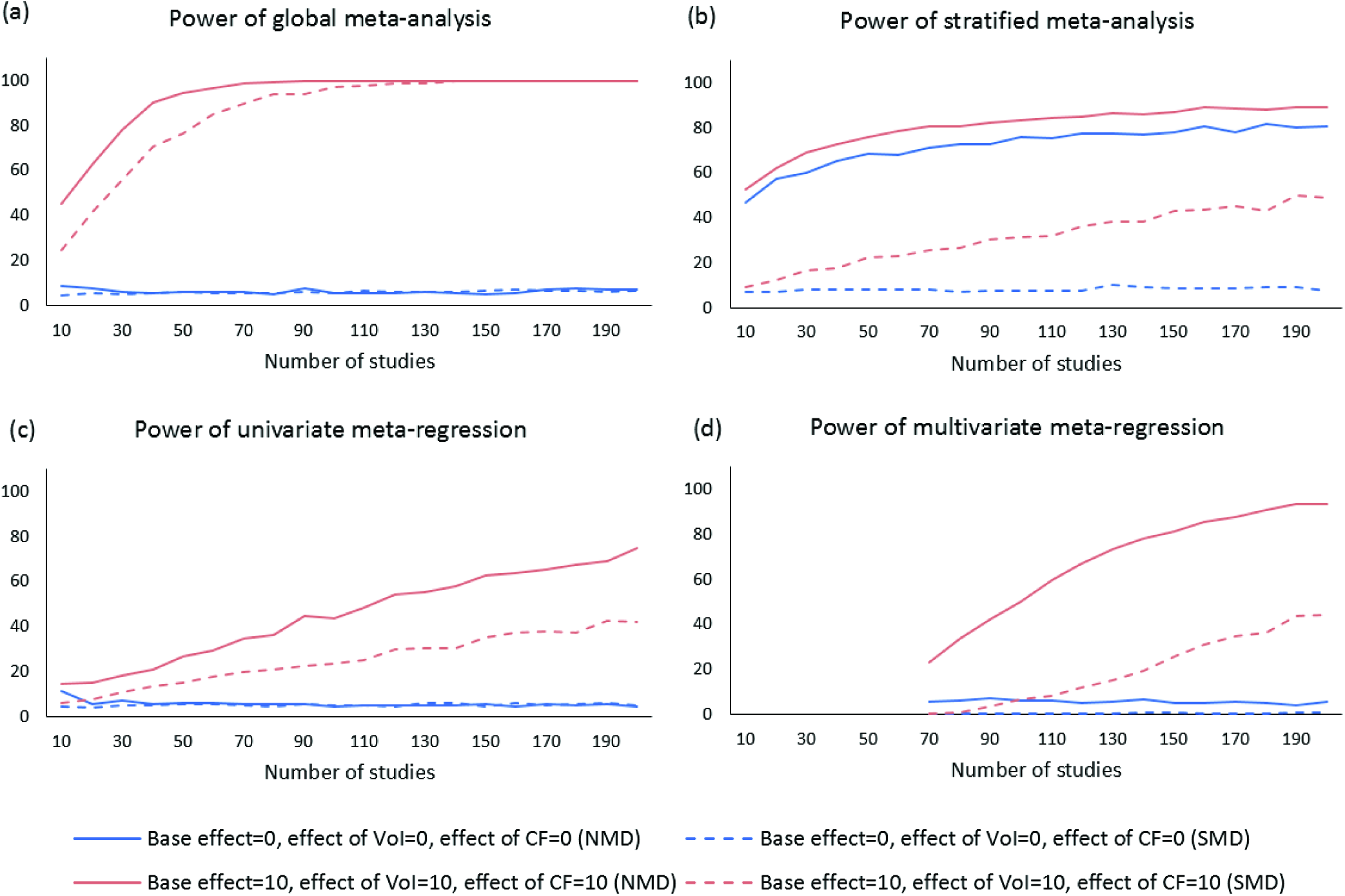
Sensitivity to the number of studies using (a) Global meta-analysis; (b) Stratified meta-analysis; (c) Univariate meta-regression; (d) Multivariate meta-regression. In each case the x-axis represents the number of studies and the y-axis represents the observed statistical power. Solid lines represent NMD estimates of effect size and the dashed lines an SMD estimate. Blue lines represent meta-analyses in which the modelled effect is zero, and red lines represent meta-analyses where an effect is present.

Next, we tested univariate meta-regression. For both NMD and SMD measures of effect size the false positive rate was constant at around 5% (Fig 1c). Power to detect the modelled difference in efficacy between subgroups increased as the number of studies increased, but was always substantially greater for NMD (44% with 100 studies, 75% with 200 studies) than with SMD (24% with 100 studies, 42% with 200 studies).

Finally, we used multivariable meta-regression. For SMD the false positive rate was around 0.4%, and with NMD estimates of effect size it was higher, but independent of study size, at around 6%. Power to detect the modelled difference in efficacy between subgroups increased as the number of studies increased, but was always substantially greater for NMD (50% with 100 studies, 94% with 200 studies) than with SMD (7% with 100 studies, 44% with 200 studies).

#### 2. Sensitivity to effect of the variable of interest

We were interested to see the effect of a variable of interest on the statistical power and the false positive rate. Figure 2 shows these findings for NMD (solid lines) and SMD (dashed lines) in the absence (blue) or presence (red) of a confounding variable of interest. As before, partitioning heterogeneity was associated with a very high false positive rate with NMD but not SMD estimates of effect size (Fig 2a). For univariate meta-regression (Fig 2b) false positive rates were around 5%, and NMD estimates of effect size had power of 44% to detect an effect of the variable of interest of 10, and around 90% to detect an effect of the variable of interest of 20. SMD estimates of effect size had power of around 24% to detect an effect of the variable of interest of 10, and around 63% to detect an effect of the variable of interest of 20. For multivariable meta-regression (Fig 2c), false positive rates were substantially lower than 5% for SMD estimates of effect size, compared with around 6% for NMD. NMD estimates of effect size had power of around 50% to detect an effect of the variable of interest of 10, and around 93% to detect an effect of the variable of interest of 20. SMD estimates of effect size had power of around 7% to detect an effect of the variable of interest of 10, and around 28% to detect an effect of the variable of interest of 20.

**Figure 2:**
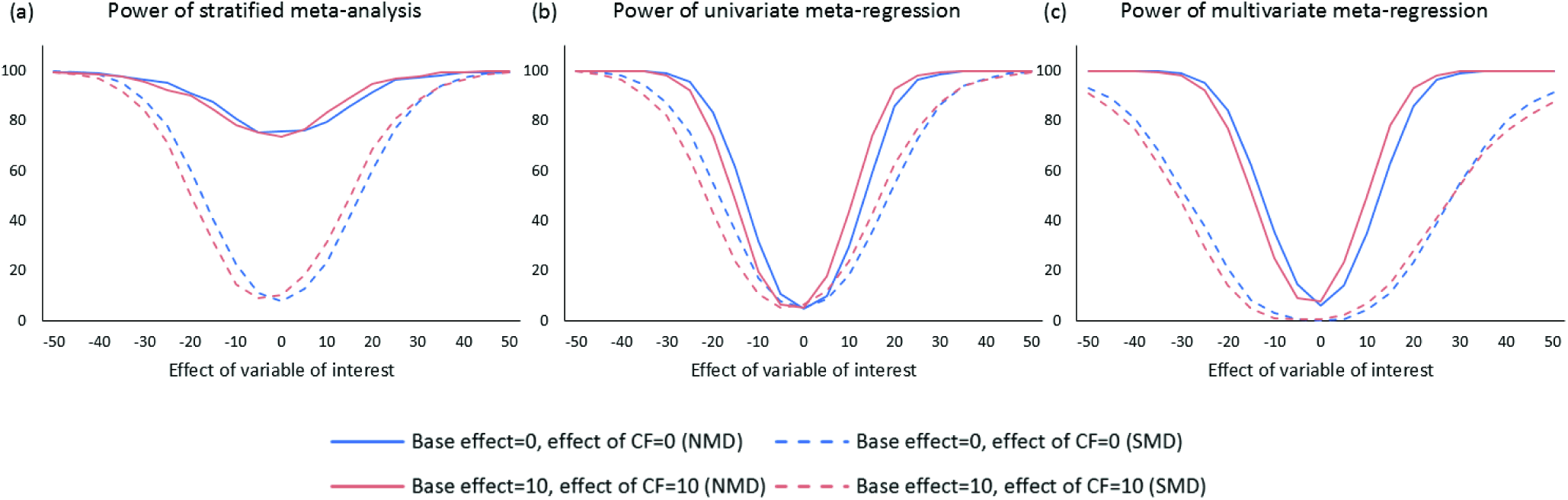
Sensitivity to the effect of the variable of interest using (a) Stratified meta-analysis; (b) Univariate meta-regression; (c) Multivariate meta-regression. In each case the x-axis represents the effect of the variable of interest and the y-axis represents the observed statistical power. Solid lines represent NMD estimates of effect size and the dashed lines an SMD estimate. Blue lines represent meta-analyses in which the modelled base and confounding variable effects are zero, and red lines represent meta-analyses where these effects are present.

#### 3. Sensitivity to effect of confounding variable

One of the main criticisms of meta-analyses of data from animal studies is that differences observed between groups of studies might not be due to the factor which defines those groups, but rather to some other, latent confounding variable which is present to a different extent in the separate groups. To establish how much of a problem this might be, we have simulated the situation where the group defining variable has no effect, but another variable asymmetrically present in the groups does have an effect. In our main scenario, based on our observations form focal ischaemia literature, the variable of interest is present in 28% of studies and the confounding variable is present in 40% of studies. We allow the representation of these to be asymmetrical such that when the variable of interest is present the prevalence of the confounding variable is 57%, and when the variable of interest is absent the prevalence of the confounding variable is 33%.

Here we now simulate the effect of varying the prevalence of the variable of interest in the presence or in the absence of the confounding variable, from 5% to 95% in increments of 5%. These are shown in Figure 3. In each case the left panel shows the proportion of tests called as being significant using an NMD estimate of effect size and the right when using an SMD estimate of effect size. The first pair of columns show results when no effect is modelled; the second when only a base effect (with no effect of the variable of interest or confounding variable) is modelled; the third when only the effect of a variable of interest is modelled, and finally when only an effect of a confounding variable is modelled. The colour represents the proportions of 1000 simulations where a statistically significant effect was reported, ranging from 0% (dark blue) to 100% (dark red); where no effect is modelled this represents the false positive rate, and where an effect was modelled this represents the statistical power.

**Figure 3:**
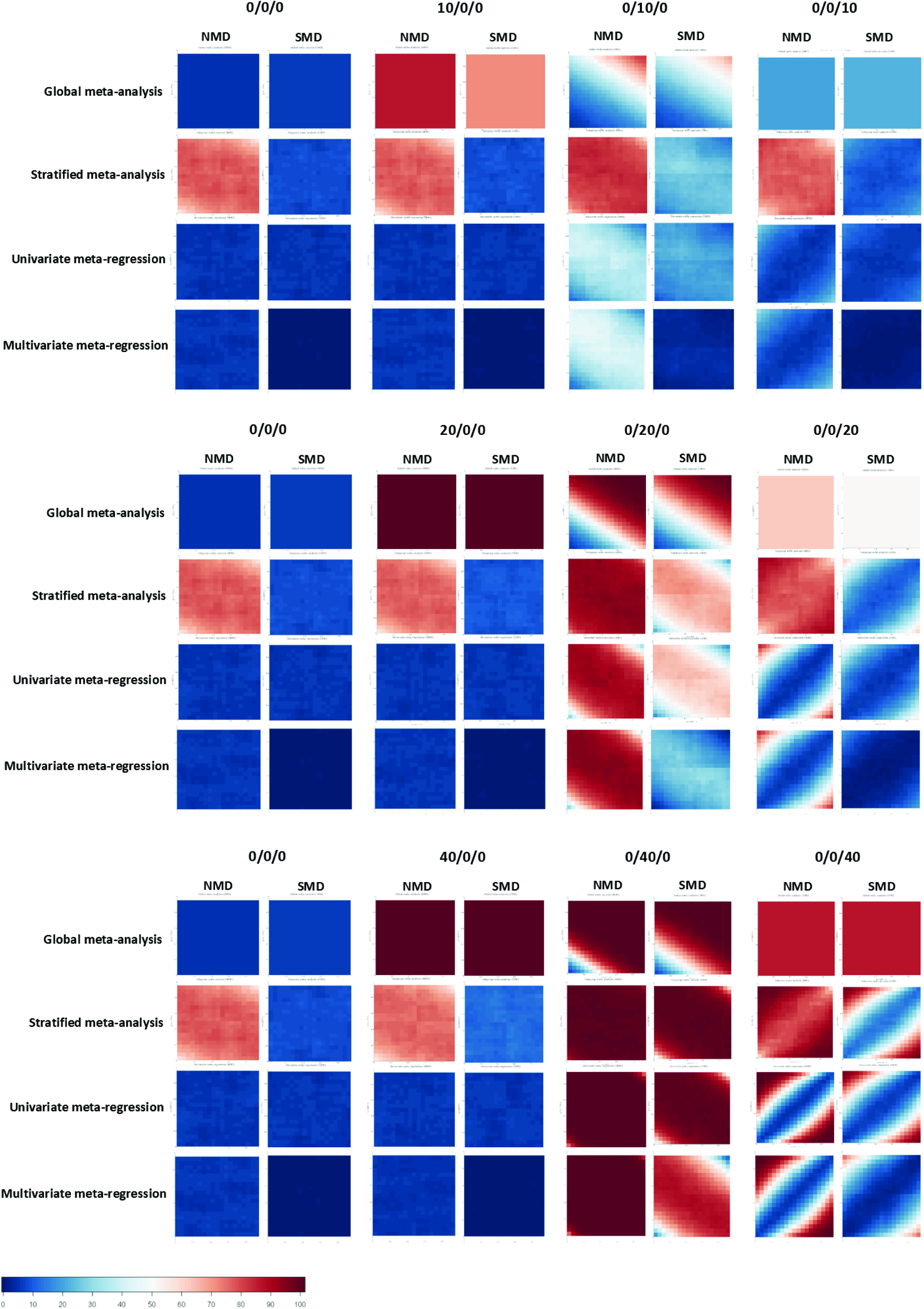
Sensitivity to the effect size of the confounding variable using NMD and SMD, and global meta-analysis, stratified meta-analysis, univariate meta-regression, and multivariate meta-regression. X/X/X represents base effect size, effect of the variable of interest and effect of a confounding variable separately. Dark blue represents zero power with lighter colours indicating increasing power up to 100% in dark red. The axes represent the prevalence of the variable of interest when a confounding variable is present (0.05 to 0.95, y-axis) or when it is absent (0.05 to 0.95, x-axis).

Each square summarises the findings of 1000 simulations each of 361 different combinations of the prevalence of the variable of interest when the confounding variable is present (0.05 to 0.95, y-axis) or when it is absent (0.05 to 0.95, x-axis). Therefore the diagonal from bottom left to top right represents the situation where the prevalence of the variable of interest is independent of the prevalence of the confounding variable; and the top left and bottom right corners represents maximum asymmetry in prevalence. We present findings for effects of the base effect, or the variable of interest, or the confounding variable, of 10 (0.1 NMD), 20 (0.2 NMD) and 40 (0.4 NMD).

As expected from the analyses above, heat maps under NMD were lighter or more close to deep red, which means the statistical power of NMD was, generally higher, compared with SMD. The false positive rate of stratified meta-analysis with NMD was particularly high (75% on average).

The heat maps show a number of other features. Firstly, when the base effect is zero but the variable of interest or the confounding variable do have an effect, the global estimate reports a significant difference, particularly as the prevalence of the variable of interest increases. Secondly, the ability of univariate and multivariable meta-regressions to detect an effect of the variable of interest are least powerful when the prevalence is very low (bottom left) or very high (top right), and most powerful when it is around 50%. Finally, the presence of a confounding variable can indeed lead to a spurious finding from univariate or multivariate meta-regression that a variable of interest is associated with differences in outcome, but for this effect to be pronounced the confounding variable needs to have a substantial impact, or the asymmetry of representation of the confounding variable in the populations defined by the variable of interest needs to be substantial, or both. Interestingly, this effect seems to be more pronounced for univariate than for multivariate meta-regression.

#### 4. Sensitivity to method of estimating tau squared

Since the method of estimating tau squared can influence the weight assigned to individual studies, it might also have an impact on the power. In the multivariate meta-regression, Hunter-Schmidt’s method and maximum-likelihood method showed a slight advantage (18.2% and 9.0% higher than the average) under the NMD method (Fig 4). For the global estimate of effect, partitioning heterogeneity and univariate meta-regression, there was not much difference among the seven methods of estimating tau squared.

**Figure 4:**
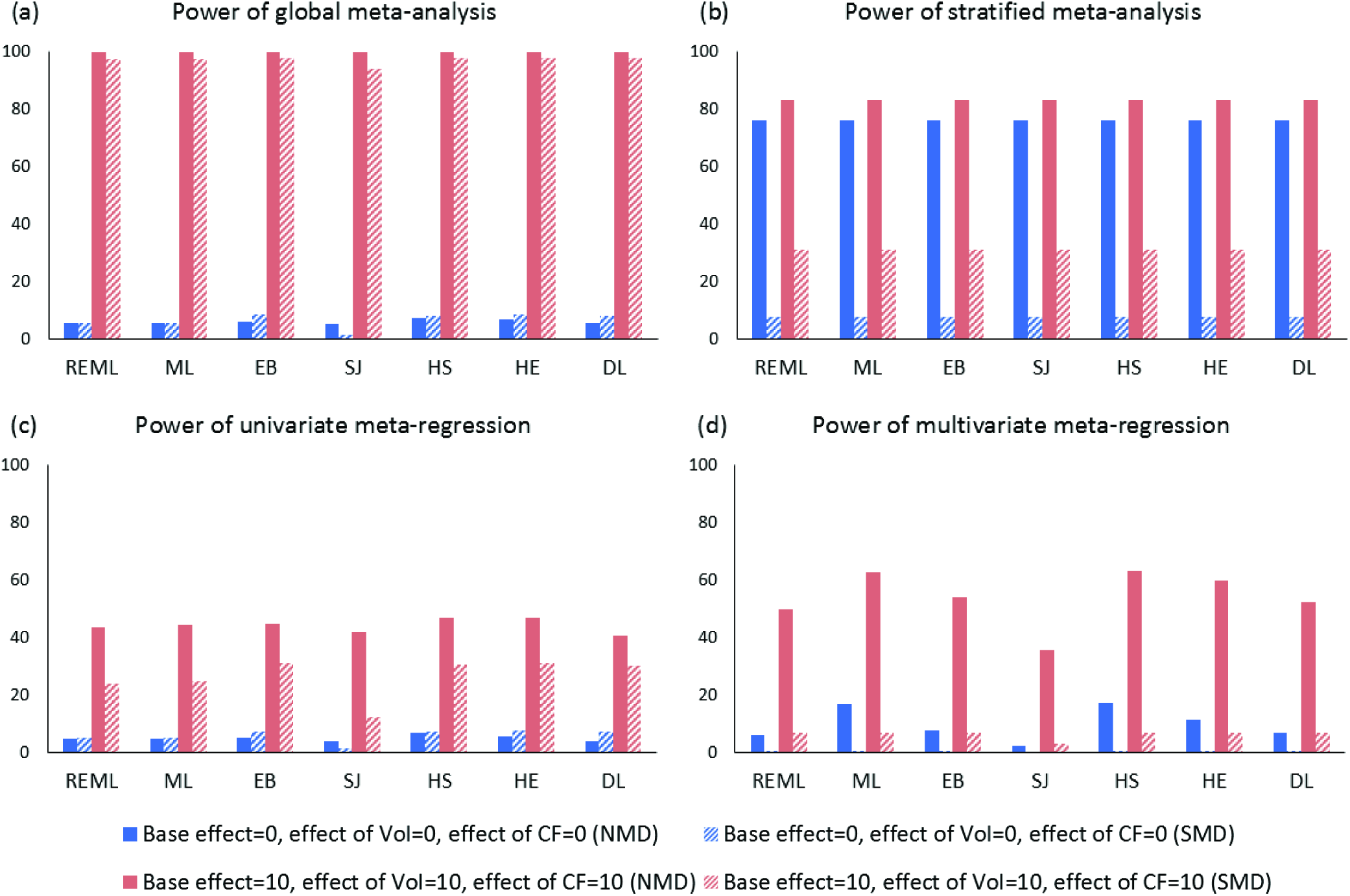
Sensitivity of the observed statistical power (y-axis) to the method of estimating tau squared (x-axis) using (a) Global meta-analysis; (b) Stratified meta-analysis; (c) Univariate meta-regression; (d) Multivariate meta-regression. REML (restricted maximum likelihood); ML (maximum likelihood); EB (Empirical Bayes); SJ (Sidik-Jonkman); HS (Hunter-Schmidt); HE (Hedges); DL (DerSimonian-Laird). Solid bars represent NMD estimates of effect size and the dashed bars an SMD estimate. Blue bars represent meta-analyses in which the modelled effect is zero, and red bars represent meta-analyses where an effect is present.

## Discussion

Our simulations provide some guidance for those conducting meta-analyses of data from animal studies in focal cerebral ischaemia. Firstly, given a choice between using NMD or SMD as an estimate of effect size, it is preferable to use the NMD approach. Where only a proportion of a dataset are amenable to such an approach a judgement should be made about the best approach, and this will depend on the number of studies that would be excluded from analysis. For global estimates of efficacy it seems that perhaps one third of studies can be lost before the power for an NMD analysis of the remainder falls below that of an SMD analysis of the entire dataset. If the main interest is in the impact of variables of interest on the observed efficacy then for both univariate and multivariable meta-analysis it appears that at least one half of studies can be lost from the NMD analysis before power falls to that obtained under SMD.

Coupled with our previous demonstration of the weakness of SMD in the ascertainment of small study effects (4) we believe that SMD measures of effect size in meta-analyses of data from animal studies should be avoided if at all possible.

Secondly, our findings suggest that the reporting of differences between sub groups on the basis of partitioning of heterogeneity is not appropriate, and investigators should use univariate or multivariate meta-regression instead. Since a series of univariate analysis will require some correction for multiplicity of testing, whereas multivariate meta-regression will not, the latter will usually be the preferred approach if sufficient studies are available. We appreciate that our advice regarding partitioning of heterogeneity runs counter to that given in the past by ourselves and others about the optimal statistical approach, but we believe that the problems with partitioning heterogeneity which we have shown are sufficient to mandate a change in practice. However, we also appreciate that many investigators will have established a statistical analysis plan in an a priori systematic review protocol, which may describe the use of NMD and partitioning of heterogeneity. We recommend that they continue with their statistical analysis plan as described, but that they also present, as a sensitivity analysis, an analysis using a meta-regression approach.

Thirdly, our data support the utility of assembling ensemble datasets of similar studies to explore the impact of aspects of study design, where it can reasonably be assumed that such factors will have similar impact across for instance different drugs being tested, or where interactions can be explored using the meta-regression approach. This is one motivation for our efforts to secure a common ontology for such datasets, that they might be made available to the community in a form that facilitates their re-use (8;9). For many drugs it may be that the totality of the research conducted to date is not able reliably to detect the effects of variables of interest, which is clearly of concern in decisions about whether there is sufficient evidence of efficacy in different circumstances to justify the decision to proceed to clinical trial. A meta-analysis with fewer than 100 contributing experiments is unlikely reliably to be able to exclude important effects of for instance gender or co-morbidity unless these effects are large.

Additionally, for the method of estimating tau squared, the iterative methods (maximum-likelihood, restricted maximum-likelihood and Empirical Bayes) did not perform substantially better than the non-iterative methods. Only for the multivariate meta-regression, Hunter-Schmidt’s method and maximum-likelihood had a slight advantage. Non-iterative methods are therefore recommended for simulations such as this, because they save substantial time when running the program; for the conduct of meta-analysis where multivariable meta-regression will be performed, we recommend either the Hunter-Schmidt or maximum likelihood approaches.

Finally, we are keen to explore the possibility of enabling this simulation tool online, so that researchers, knowing the number of studies likely to be available to them, can decide what is the most efficient and informative statistical approach for their data. Our code is available on GitHub but has not yet been optimised for a parallel computing environment, and we are actively exploring the possibility of hosting the code on a server with sufficient processing power to render the simulations in a timely fashion.

Our approach has some weaknesses. Firstly, because our simulation was derived from studies in focal cerebral ischaemia, it is not clear whether they will also be relevant to meta-analyses of other disease models. It is our experience that datasets collected in systematic reviews of findings from animal studies modelling Parkinson’s disease (10), Alzheimer’s disease (11), multiple sclerosis (12), spinal cord injury (13), and glioma (14) are broadly similar, and we believe the same problems may arise. Where the number of individual studies is smaller and those studies are individually larger (as with meta-analyses of human studies), we do not know if this is important. Secondly, the coefficients which we derived from real data to build the regression model (used for calculating true effect size and generating mean and standard deviation for simulated studies) were derived using the NMD approach. To compare the approaches we had to use the same model, and since our variables were selected from the NMD model this may have advantaged this approach. Thirdly, we treated two continuous variables as discrete variables. The median of each group was taken as a representative value to aid computation, but it may have influenced the precision of our approach. Finally, our simulations have explored the effect of a binary variable of interest (i.e. the presence or absence of a factor); we have not explored performance when the variable of interest can take more than 2 values.

